# INTEGRATE: Model-based multi-omics data integration to characterize multi-level metabolic regulation

**DOI:** 10.1101/2021.08.13.456220

**Authors:** Marzia Di Filippo, Dario Pescini, Bruno Giovanni Galuzzi, Marcella Bonanomi, Daniela Gaglio, Eleonora Mangano, Clarissa Consolandi, Lilia Alberghina, Marco Vanoni, Chiara Damiani

## Abstract

Metabolism is directly and indirectly fine-tuned by a complex web of interacting regulatory mechanisms that fall into two major classes. First, metabolic regulation controls metabolic fluxes (i.e., the rate of individual metabolic reactions) through the interactions of metabolites (substrates, cofactors, allosteric modulators) with the responsible enzyme. A second regulatory layer sets the maximal theoretical level for each enzyme-controlled reaction by controlling the expression level of the catalyzing enzyme. In isolation, high-throughput data, such as metabolomics and transcriptomics data do not allow for accurate characterization of the hierarchical regulation of metabolism outlined above. Hence, they must be integrated in order to disassemble the interdependence between different regulatory layers controlling metabolism. To this aim, we proposes INTEGRATE, a computational pipeline that integrates metabolomics (intracellular and optionally extracellular) and transcriptomics data, using constraint-based stoichiometric metabolic models as a scaffold. We compute differential reaction expression from transcriptomic data and use constraint-based modeling to predict if the differential expression of metabolic enzymes directly originates differences in metabolic fluxes. In parallel, we use metabolomics to predict how differences in substrate availability translate into differences in metabolic fluxes. We discriminate fluxes regulated at the metabolic and/or gene expression level by intersecting these two output datasets. We demonstrate the pipeline using a set of immortalized normal and cancer breast cell lines. In a clinical setting, knowing the regulatory level at which a given metabolic reaction is controlled will be valuable to inform targeted, truly personalized therapies in cancer patients.

**Author summary:** The study of metabolism and its regulation finds increasing application in various fields, including biotransformations, wellness, and health. Metabolism can be studied using post-genomic technologies, notably transcriptomics and metabolomics, that provide snapshots of transcripts and metabolites in specific physio-pathological conditions. In the health field, the transcriptome and, more recently, the metabolome have been broadly profiled at the pre-clinical and clinical levels. The informative power of single omic technologies is inadequate since metabolism regulation involves a complex interplay of regulatory steps. While gene expression regulates metabolism by setting the upper level of metabolic enzymes, the interaction of metabolites with metabolic enzymes directly auto-regulates metabolism. Therefore there is a need for methods that integrate multiple data sources. We present INTEGRATE, a computational pipeline that captures dynamic features from the static snapshots provided by transcriptomic and metabolomic data. Through integration in a steady-state metabolic model, the pipeline predicts which reactions are controlled purely by metabolic control rather than by gene expression or a combination of the two. This knowledge is crucial in a clinical setting to develop personalized therapies in patients of multifactorial diseases, such as cancer. Besides cancer, INTEGRATE can be applied to different fields in which metabolism plays a driving role.

## Introduction

In recent years, the biological role of metabolism has been reconsidered. Being closely integrated with most - if not all – cellular processes, metabolism may act as a sensitive integrative readout of the physio-pathological state of a cell or organism [1]. Consistently, many physio-pathological states and multifactorial diseases, from cancer to neurodegeneration and aging, show a specific metabolic component [2].

While the general topology of metabolism is well established, the characterization and understanding of system-level regulation of metabolism remain largely unresolved, although some general rules emerged in recent years [3, 4]. Each metabolic flux depends on metabolic enzymes, whose levels and catalytic activities are the outcome of gene expression and regulatory events that include epigenetic control of chromatin, accessibility to transcription factors, the rate of transcription of the individual genes encoding metabolic enzymes, as well as post-transcriptional and post-translational events (from RNA splicing to enzyme phosphorylation). These complex, hierarchical regulatory mechanisms are orchestrated by signal transduction pathways and set the upper level for the flux of each enzyme-catalyzed metabolic reaction. An increasing body of evidence [5–7] indicates that metabolism is not passively regulated by the hierarchical control mechanisms outlined above: on the contrary, metabolism auto-regulates metabolic fluxes (i.e., the rate of individual metabolic reactions) through the interactions of metabolites (substrates, cofactors, allosteric modulators) with the responsible enzymes. An enzyme reaction rate depends on the concentration of its substrate(s) unless the substrate is saturating the enzyme (i.e., is in significant excess over the enzyme Michaelis-Menten constant *K_M_*). The reaction substrate and other metabolites within the same pathway or belonging to other cross-related biochemical pathways can fine-tune each enzyme-catalyzed reaction through allosteric effects that effectively up- or down-modulate the ability of the enzyme to catalyze the reaction at a given substrate concentration. Since substrates and allosteric effectors are consumed and produced by other metabolic reactions and exchanged with the extracellular environment, this metabolic control layer contributes to regulating metabolism at the system level and may even fully account for metabolic rewiring [8]. It is worth noting that the hierarchical expression and the metabolic control layers are not entirely independent. The gene expression layer sets the upper bound for any given flux, and metabolites control the gene expression cascade at different levels, from epigenetic modification of chromatin [9] to signal transduction [10], from enzyme phosphorylation [10] to transcription [11].

Hence, differences in metabolic fluxes are only partially determined by variation in protein/gene expression. Let us take, for example, two cells A and B at steady-state and a specific, irreversible metabolic reaction *r*_1_ (*S*_1_ → *P*_1_) catalyzed by enzyme *E*_1_. Significantly higher activity in cell A of enzyme *E*_1_, via *transcriptional regulation*, might originate - or not - a change of flux through reaction *r*_1_, according to the following general cases:

- *trascriptional control*: *S*_1_ is in excess, so the flux through *E*_1_ is independent of [S1] and depends only on the level of *E*_1_. The supply of S1 from the environment or other network reactions should increase to keep up with the higher consumption rates of S1, avoiding a decrease of [*S*_1_] below the level where the reaction does not depend on [*S*_1_];
- *metabolic control*: E1 is in excess, so the flux through E1 is independent of its level and depends only on changes in [S1];
- *metabolic and transcriptional control*: the flux through E1 is co-regulated with changes in [S1].

Characterizing the landscape of metabolism and its regulation is of paramount importance in various fields, including health, wellness, and biotransformations [12]. The first requirement for this characterization is the knowledge of metabolic fluxes. However, direct determination of metabolic fluxes through the use of labeled substrates lags behind other omic technologies, such as metabolomics and transcriptomics, mainly due to technical difficulties [13], especially at the sub-cellular level [14]. On the contrary, transcriptomics (or proteomics) and metabolomics datasets are increasingly being collected in large cohorts but do not allow for accurate characterization of the regulatory mechanisms controlling metabolism unless opportunely integrated. More recently, parallel transcriptomic and metabolomic datasets started to appear [15–20]. Yet, the integration of these different omic data has so far been generally limited to gene-metabolite correlation analysis or pathway enrichment analysis of genes and metabolites [21,22]. Hence, there is a need for data science methods to integrate transcriptomic and metabolomic data to capture all the facets of the interdependence between metabolism and gene expression.

Constraint-based steady-state models represent a valuable framework to predict metabolic fluxes from the other high-throughput omics data. In particular, a plethora of methods have been conceived to integrate transcriptomic data into these kinds of models by relying on Gene-Protein-Reaction associations (GPRs) encoded within them, as reviewed in [23–25]. Intracellular metabolomics data have also been indirectly integrated into constraint-based steady-state models in the form of constraints on fluxes [26–28], aiming at identifying the metabolic flux distribution better fitting the given data.

Current model-based attempts to discern trascriptionally from metabolically controlled fluxes present some limitations [29, 30]. Katzir et al. [30] do not directly use metabolomics data to infers reactions controlled at the metabolic level using metabolomic data but determine them by elimination, as fluxes that are not regulated at the transcriptional, translational, or post-translational level. On the contrary, the approach by Cakir at al. [29] is based on the concept of neighborhood in a graph, it does not distinguish reactions substrates from products, nor enzyme subunits from isoforms, and does not predict whether a reaction is up or down-regulated, but simply de-regulated.

We here present the INTEGRATE (Model-based multi-omics data INTEGRAtion to characterize mulTi-level mEtabolic regulation) pipeline to accurately characterize the landscape of metabolic regulation in different biological samples, starting form metabolomics data (and optionally exometabolomic data to derive utilization and elimination of selected nutrients and waste products) and transcriptomic data. INTEGRATE first computes differential expression of reactions from transcriptomics data (transcriptional regulation only). Then, it exploits constraint-based modeling to predict how the global relative differences in expression are expected to translate into consistent differences in metabolic fluxes. To improve model predictions, INTEGRATE optionally sets constraints also on selected extracellular fluxes, according to exo-metabolomic data. In parallel, INTEGRATE uses intracellular metabolomic datasets and the mass action law formulation to predict how differences in substrate availability translate into differences in metabolic fluxes (metabolic regulation only), neglecting enzymatic activity. The intersection of the two output datasets discriminates fluxes regulated at the metabolic and/or gene expression level.

A a proof-of-principle of the pipeline, we used a set of immortalized normal and cancer breast cell lines and reconstructed a manually curated, multi-compartment metabolic network of human central carbon metabolism.

## Results

### The INTEGRATE pipeline

We conceived the INTEGRATE methodology to particularize the hierarchical regulation of metabolic differences across different groups of biological samples. For the sake of readability, we refer to different groups of samples simply as cell lines, which represent the application scope of this work, bearing in mind that the methodology is general and can be applied to any group of samples, as for instance, tissues from different patients.

INTEGRATE takes as input 1) a generic metabolic network model, including GPRs 2) transcriptomics data 3) intracellular metabolomics data 4) extracellular fluxes data.

INTEGRATE returns as final main output two list of metabolic fluxes: 1) fluxes that vary across cells consistently with both metabolic and transcriptional regulation 2) fluxes that vary consistently with metabolic regulation only.

The core process of INTEGRATE methodology is depicted in Figure 1 and consists of integrating the input experimental datasets, which are centered around heterogeneous objects (i.e., genes, metabolites and fluxes) into the input metabolic network in order to obtain the three following datasets, each of which is centered around the object reaction:

*Reaction Activity Scores (RAS)*. This dataset includes for each input model reaction and for each sample a RAS score. The score is based on the expression value (RNA-seq read counts) of the genes encoding for catalyzing enzymes and on the relationship among them, as previously done in [31].
*Feasible Flux Distributions (FFD)*. This dataset includes a large number of flux distributions associated with a given cell line, obtained by uniformly sampling the feasible flux region of the metabolic model. The model has previously been tailored on the cell line by integrating transcriptomics and relative extracellular constraints.
*Reaction Propensity scores (RPS)*. This dataset includes for (ideally) each input model reaction and for each sample, a RPS score based on the availability of reaction substrates. The score is computed as the product of the concentrations of the reacting substances, with each concentration raised to a power equal to its stoichiometric coefficient. According to the mass action law, the rate of any chemical reaction is indeed proportional to this product. This assumption holds as long as the substrate is in significant excess over the enzyme constant *K_M_*. If one single reaction substrate is missing in the metabolomics measurements, the reaction is omitted from the dataset.

**Fig 1.**
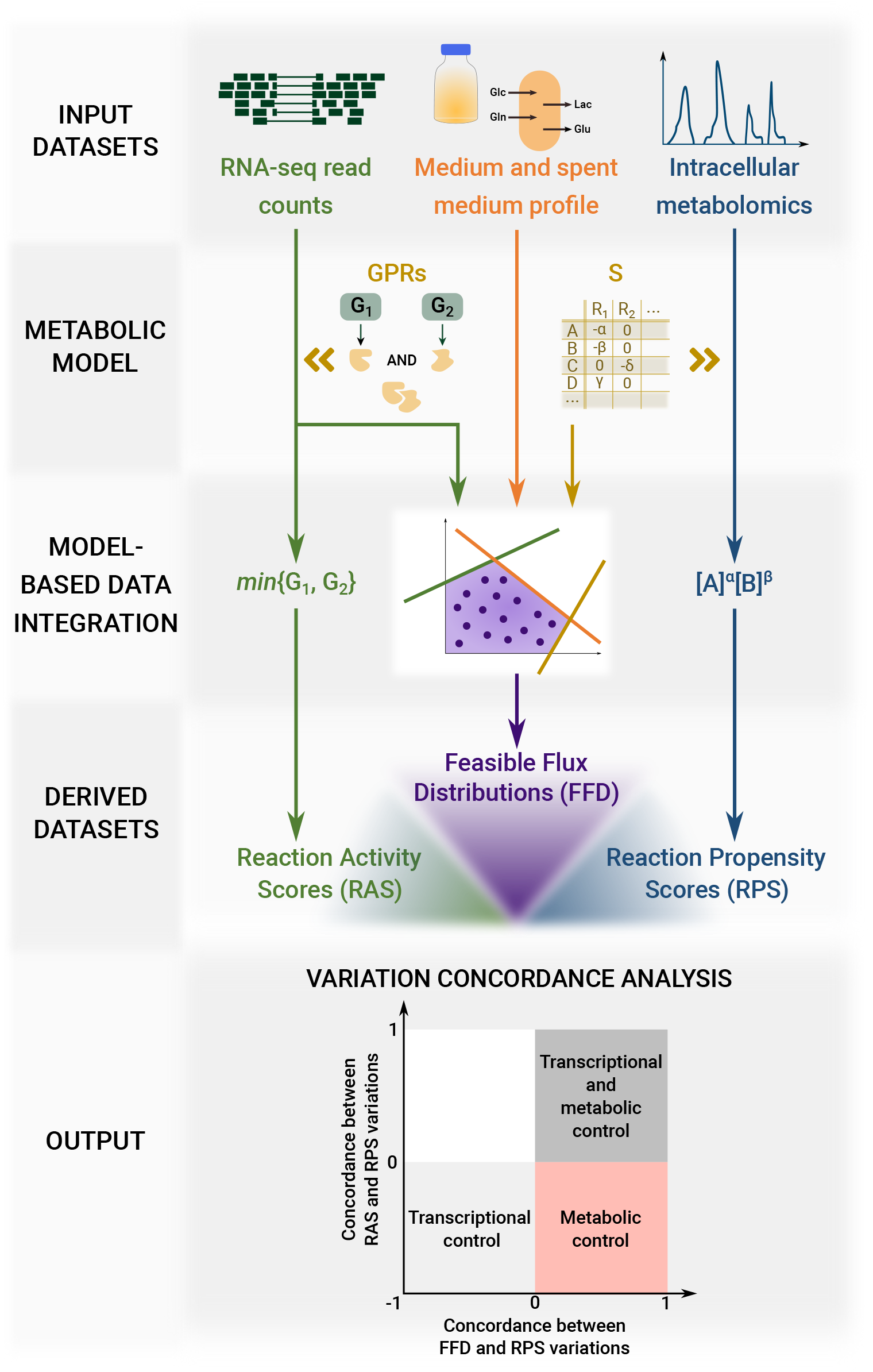
Graphical representation of INTEGRATE pipeline. Inputs, intermediate and final outputs are reported.

Once the three reaction-oriented datasets are obtained, INTEGRATE assesses for each of them whether the value of each reaction is significantly higher or lower in a given cell line as compared to another one. We consider a variation as statistically significant if both the null hypothesis is rejected according to a proper statistical test and if the variation is greater than a threshold value.

INTEGRATE then assigns two scores to metabolic reactions. The first score quantifies the concordance level between the variation signs obtained for the RAS dataset and those obtained for the RPS dataset (for reactions in common). Highly concordant reactions correspond to fluxes whose metabolic and transcriptomic regulation is concerted, poorly concordant vice versa. The second score assesses the concordance between FFD and RPS (for reactions in common) and thus whether flux variations are consistent with metabolic regulation. Reaction displaying a low RAS-RPS agreement but a high FFD-RPS correspond to *metabolically controlled* reactions.

We remark that we preferred not to give the same attention to flux variations that are consistent with transcriptional regulation only, based on the concordance between RAS and FFD, because the two datasets are not independent.

Scripts to reproduce the overall workflow are available at: https://github.com/qLSLab/integrate.

### Selected breast cancer cell lines display heterogeneous metabolic profiles at balanced growth

To test our approach, we applied INTEGRATE to cell lines that we expected to be metabolically heterogeneous. We selected four breast cancer cell lines deriving from either primary of metastasis breast cancer tissues belonging to different molecular classifications, and a non-tumorigenic breast cell line. The name, origin and molecular subtyping of the five cell lines are summarized in Table 1.

**Table 1.** Characterization of the investigated non-tumorigenic and cancer breast cell line in terms of their subtype and origin.

| Cell line | ER | PR | Subtype |  | Origin |
| --- | --- | --- | --- | --- | --- |
|  |  |  | HER2 | Luminal |  |
| MCF102A |  |  |  |  | Non-tumorigenic breast epithelial cell line |
| SKBR3 | - | - | + |  | Breast adenocarcinoma |
| MCF7 | + | + | - | A | Pleural effusion metastasis of a breast adenocarcinoma |
| MDAMB231 | - | - | - |  | Pleural effusion metastasis of a breast adenocarcinoma |
| MDAMB361 | + | + | + | B | Non-tumorigenic breast epithelial cell line |

The five cell lines were cultured in a similar growth medium. It can be observed in Figure 2A that the cell lines present major differences in terms of proliferation rate.

**Fig 2.**
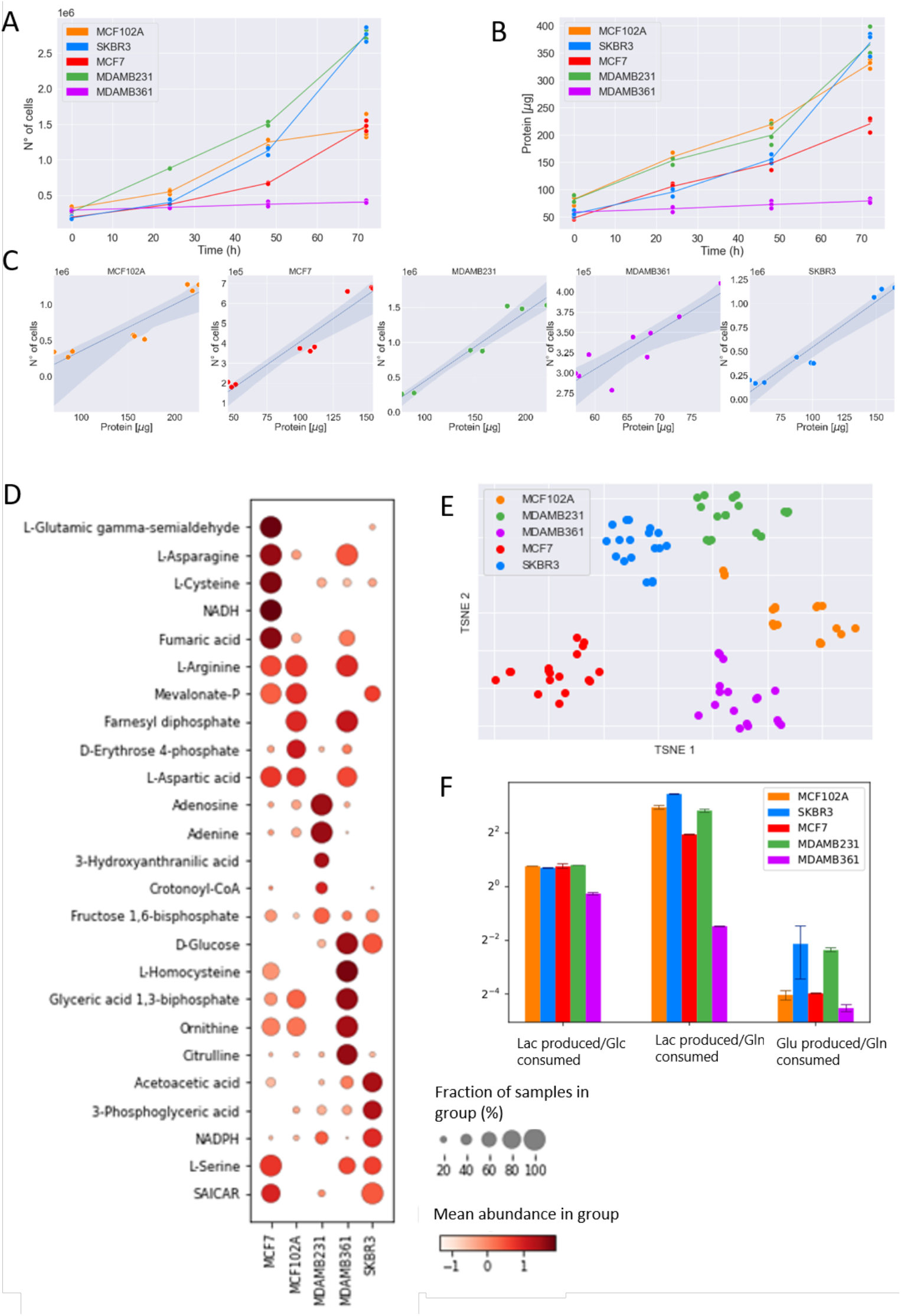
Experimental metabolic measurements at balanced growth phase. A) Number of cells in time. B) Protein content in time. C) Correlation plots between protein content and number of cells for experimental observations in the 0-48hrs time windows. D) Dotplot representing the mean metabolite abundances within each line (visualized by color) and the fraction of samples of each cell line with the abundance over the average of all cell lines samples (visualized by the size of the dot). The first five metabolites that better distinguish each cell line from the others, according to t-test p-value, are reported. E) t-SNE dimensionality reduction of intracellular metabolomic profiles. F) Extracellular flux ratios, derived from spent medium measurements.

We first identified a balanced growth phase suitable to obtain measurements to be used as constraints in steady state modeling. To this aim, we analyzed both the number of cells in time and the protein content (Figures 2A-B). It can be observed in Figure 2C that, between 0 and 48 hours, the protein content linearly correlates with the number of cells, indicating that the cell size is constant in this time window. We, therefore, concentrated further analyses on the 0-48 hours time window.

We estimated the consumption and production rates of lactate, glutamine, glucose and glutamate in the 0-48 hours time interval from YSI analysis of spent medium. We focused on these metabolites because glucose and glutamine are the main carbon sources of cancer cells and because the rate of lactate and glutamate production over glucose and glutamine consumption is notoriously difficult to be properly predicted by constraint-based models [32, 33]. We quantified the abundance of intracellular metabolites and prepared libraries for RNA-sequencing at 48h.

It can be observed in Figure 2E that the cell lines present major differences in terms of metabolic profile. The clusters of samples referring to each cell are indeed well separated from one another. In particular, as it can be observed in the dot plot in Figure 2D, the abundance of some metabolites well distinguish a cell from the others. MCF7 cell line shows a prevalence of NADH involved in redox balance, and fumaric acid that is involved in the tricarboxylic acid cycle. MCF102A is enriched in the pentose phosphate pathway as D-Erythose 4-phosphate. MDAMB231 cell line results enriched in metabolites involved in methionine and cysteine metabolism (Adenosine) and nucleotide synthesis (Adenine). MDAMB361 cell line shows enrichment in metabolites involved in glycolysis (D-glucose and Glyceric acid 1,3-biphosphate), methionine metabolism (L-homocysteine) and urea cycle (ornithine and citrulline). Finally, the most abundant metabolites in SKBR3 cell line are involved in glycolysis (D-glucose and 3-Phosphoglyceric acid), cholesterol synthesis (Acetoacetic acid) and redox balance (NADPH).

Measurements on spent medium in Figure 2F show that the ratio of lactate produced over glucose consumed is quite similar across the five cell lines. In contrast, the lactate and glutamate over glucose ratios are more heterogeneous.

All raw experimental data are provided in Supplementary File 1.

### The ENGRO2 metabolic model

Genome-wide metabolic reconstruction of human metabolism, such as Recon3D [34] are precious repositories of detailed and multi-level information about human metabolism. They involve thousands of metabolites, reactions and genes. In principle, they can be used directly in our pipeline as a scaffold model for integrating the experimental input data relative to the five breast cell lines. Yet, in their current form, their comprehensiveness comes at the cost of some simulation issues and difficulties in interpretation of the outcomes. The most relevant simulation problem is the presence of thermodynamically infeasible loops, which make the simulated growth rate insensitive to variation in essential nutrient availability constraints, as reported in [35]. For this reason, we preferred to rely on a core model, extracted from Recon3D, focusing on more limited aspects of metabolism, but that underwent extensive manual curation and debugging.

To this aim, we reconstructed the ENGRO2 metabolic network, which is a constraint-based core model about central carbon metabolism and essential amino acids metabolism. ENGRO2 is a follow-up of the core model of human central metabolism ENGRO1 introduced in [36] to gain new knowledge about the logic of metabolic reprogramming in promoting tumour cells proliferation under different nutritional conditions.

Starting from ENGRO1, we reconstructed a more extended and curated constraint-based core model of human metabolism. The reconstruction of the ENGRO2 model was based on a step-wise manual procedure using ENGRO1 model as a scaffold and progressively including specific pathways or reactions from Recon3D according to their relevance in literature for cancer cells. In order to associate Gene-Protein-Reaction associations, we relied, when possible, on the HMR core model [37], whose GPRs were manually refined, and for remaining reactions on the RECON3D model [31].

The most invasive change we implemented in the model was the compartmentalization of reactions and metabolites within the intracellular space. Many studies support the evidence of an altered expression of some mitochondrial carriers in multiple cancer cells that, most probably, arise as an adaptation to their current metabolic state and the consequent new requirements [38]. Therefore, in addition to the extracellular compartment, we divided the internal model side into cytosol and mitochondrion, and we catalogued all the included reactions depending on their localization according to literature knowledge. This first change implied the necessity to add specific transport reactions in order to link biochemical transformations occurring within the cytosol and mitochondrial matrix. The extension of ENGRO1 to the ENGRO2 model also included the addition of all the reactions belonging to non-essential and essential amino acids metabolism. Further details on the modifications with respect to ENGRO1 are reported in the Methods section.

The reconstructed model has been refined by checking: i) the capability to reproduce ENGRO1 results [36] in terms of contribution of glucose and glutamine as carbon and nitrogen sources for supporting proliferative wirings and of sensitivity of the model at high and low levels of these nutrients; ii) the actual essentiality of essential amino acids, that is, null growth rate if they are depleted from the medium; iii) the capability to reproduce experiments in literature, including the dependence of cancer phenotypes from the *de novo* synthesis of palmitate-derived lipids rather than on an external source of fatty acids, as came out in [39], the *in silico* simulation of the effect of an inverse agonist for the nuclear receptor liver-X-receptor, whose role is to regulate the expression of some key genes in the glycolysis and lipogenesis, as a putative cancer treatment approach [40], the role of the creatine kinase (CK) enzyme that due to the requested high amount of ATP may act as potential anticancer agent [41], and the overexpression of argininosuccinate synthase enzyme as a mechanism for impairing cancer cells proliferation due to aspartate deviation from the production of pyrimidines [42].

The final version of the ENGRO2 core model consists of 496 reactions and 422 metabolites. A graphical representation of the network, splitted into two figures for improved readability, is reported in Supplementary Figure 1 and 2, depicting, respectively, the central carbon metabolism and the metabolism of the essential amino acids. The model in SBML format is provided as Supplementary File 2.

### Cell-relative metabolic models

We customized the ENGRO2 core model to obtain five cell-specific core models of the cell lines under study. Because the cell-specific models must be functional to highlight the metabolic differences between the cell lines, we incorporated most constraints in the form of relative constraints. For this reason, the models cannot be considered as cell-specific stand-alone models. Hence we refer to them as *cell-relative*.

Specifically, we integrated the following three kind of relative constraints:

1. *Constraints on nutrient availability*. Only metabolites that are supplied in the medium can be internalized by the network. These constraints also reflect the slight differences among the growth medium of the five cells. Similarly to what was done in [43], if the concentration of metabolite X in the medium of cell A is, for instance, 20% higher than in the medium of Cell B, the maximum uptake flux allowed for metabolite X in cell A will be 20% greater than that of cell B.
2. *Constraints on extracellular fluxes*. For the metabolites for which we have estimated the consumption and production rate, we set constraints on their ratios, namely on the glutamate/glutamine, lactate/glucose and lactate/glutamine ratios. The choice of constraining the relative ratios among these metabolites rather than their absolute intake or secretion rates is motivated by the limited subset of measured metabolites and the need to avoid an imbalance between such values and the arbitrary absolute values of type 1 constraints. In this way, the relative ratios between boundaries on extracellular fluxes are preserved both within and across cells.
3. *Transcriptomics-derived constraints on internal fluxes*. Each reaction is assigned a Reaction Activity Score (RAS), according to the expression of its associated genes and on the relationship among them encoded within GPR associations. The differences in the flux boundaries of any given reaction across the five cell lines reflect the differences in their RAS. More in detail, the cell line with the highest RAS is allowed to reach the maximum possible flux value, which in turn is determined by type 1 and type 2 constraints, whereas the other cell lines have a RAS-proportionally reduced capability.

We expected the integration of the above constraints to segregate the feasible flux distributions of each cell line. To verify this hypothesis, we extensively sampled the feasible region of each model. We then applied a t-distributed stochastic neighbor embedding (t-SNE) algorithm to visualize the high-dimensional sampled flux distributions location in a two-dimensional space [44].

In Figure 3 it is evident how the iterative application of the three constraints progressively well separates the flux distributions sampled from each model (corresponding to the specific colour in the plots) from one another. Notably, constraints on extracellular fluxes alone do not allow to discriminate the feasible flux distributions of the five models. On the contrary transcriptomics-derived constraints alone result in a good separation of the feasible regions of the five models. Combination of both kind of constraints decrease the distance between the sampled solutions for a given model (intra-model) and increase inter-model distances. The same conclusions are derived by also considering the correlation between the growth yield on glucose computed starting from wet and computational data (on average over the sampled FFDs). The wet growth yield is computed for each of the two collected biological replicates at 48 hours as the ratio of the protein content over the glucose consumption deriving from the YSI analyzer of spent medium. The counterpart *in silico* growth yield is computed as the mean ratio of the biomass synthesis over the the glucose uptake flux values. The condition where transcriptomics-derived constraints alone are integrated well discriminates the five cell lines in terms of their growth rate, as shown in the relative correlation plot in Figure 3. However, predictions of growth rates improve when constraints on extracellular fluxes are also added. Note that YSI-derived constraints alone are sufficient to well predict the growth rates, but they are not enough to discriminate the FFD of the five models.

**Fig 3.**
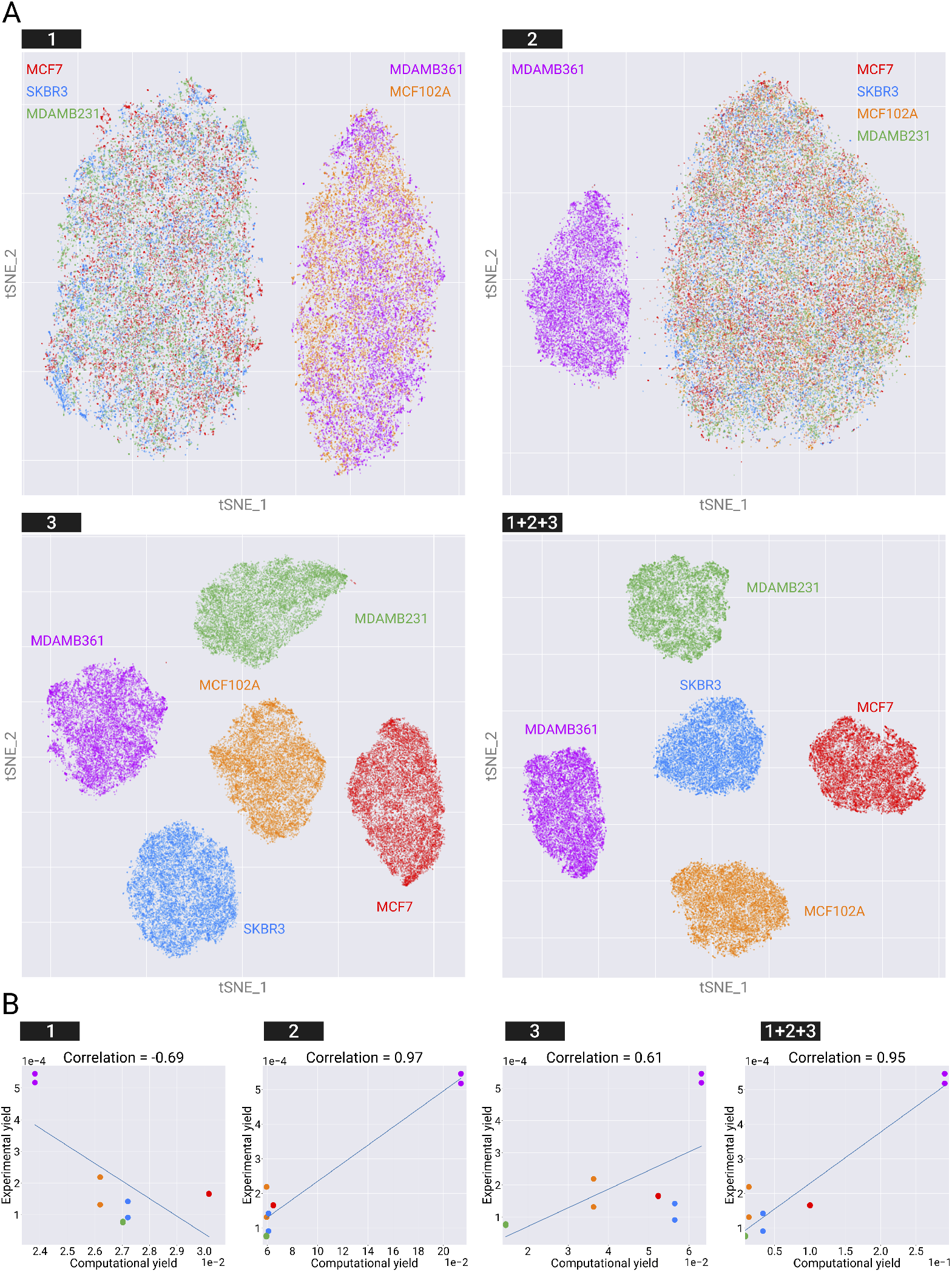
Evaluation of the iterative integration of constraints in the ENGRO2 model. A) Effect of constraints on nutrients availability (type 1), extracellular fluxes (type 2) and intracellular fluxes based on transcriptomics data (type 3) in segregating the five investigated cell lines, when they are considered alone and all together (type 1+2+3). A two dimensional map of the FFDs of the five cell line in each setting is shown. For reversible reactions, that net flux is considered. For computational reasons, only a randomly selected subset of 10000 – out of the 1 million steady-state solutions that we sampled within the feasible region of each model – is plotted. B) Correlation between the wet and *in silico* growth yield on glucose is reported for each of the four settings in panel A. The Pearson correlation coefficient is reported on top of each plot.

The final five cell-relative metabolic models are included in Supplementary File 3.

### INTEGRATE discriminates reactions regulated at different levels

By integrating the information of the three derived datasets, we can ascertain at which level each reaction is controlled. We measured the qualitative concordance between the RAS and RPS, as well as between RPS and FFD values in all pairwise comparisons between cell lines, for all eligible reactions, i.e., reactions for which metabolite levels for all substrates are available. We focused on the qualitative concordance, that is, the concordance of variation sign, rather than on a quantitative concordance of numerical variations values, as we cannot expect proportionality between RPS and FFD, nor between RAS and RPS values.

Qualitative concordance was measured by the Cohen’s Kappa coefficient, which quantifies the difference between the rate of agreement that is actually observed and the rate of agreement that would be expected purely by chance. The value of Cohen’s kappa is 1 if the two datasets are fully concordant; 0 if they agreed only as often as they would by chance. A negative value of Cohen’s kappa indicates that the two datasets agreed even less often than they would by chance. A value of –1 means that the two raters made opposite judgments in every case. This metric allowed us to rank reactions according to their concordance. We remark that Cohen’s kappa is not a statistical test that provides a well defined yes/no result. However, it has been recommended [45] to consider a value below 0.2 as poor concordance, a value between 0.21 and 0.40 as fair, between 0.41 and 0.60 as moderate, between 0.61 and 0.80 as good, and between 0.81 and 1.0 as very good agreement.

Figure 4A reports the concordance level between RAS and RPS variations (briefly RPSvsRAS) versus the concordance level between RPS and FFD variation (briefly RPSvsFFD), for the 81 metabolic reactions of ENGRO2 for which quantification of all substrate abundances was available. It can be observed that reactions distribute among the following categories:

- Reactions displaying positive values for both RPSvsFFD and RPSvsRAS scores (first quadrant - gray shadow in Figure 4A). Variations in these reactions must be imputed to transcriptional and metabolic regulation.
- Reactions having positive values for RPSvsFFD and negative values RPSvsRAS scores (fourth quadrant - pink shadow in Figure 4A). Variations in these reactions must be imputed to metabolic control only.
- Reactions having negative values for both RPSvsFFD and RPSvsRAS scores, but high values for RASvsFFD concordance (third quadrant - white shadow in Figure 4A). Variations in these reactions must be imputed to transcriptional control only.

**Fig 4.**
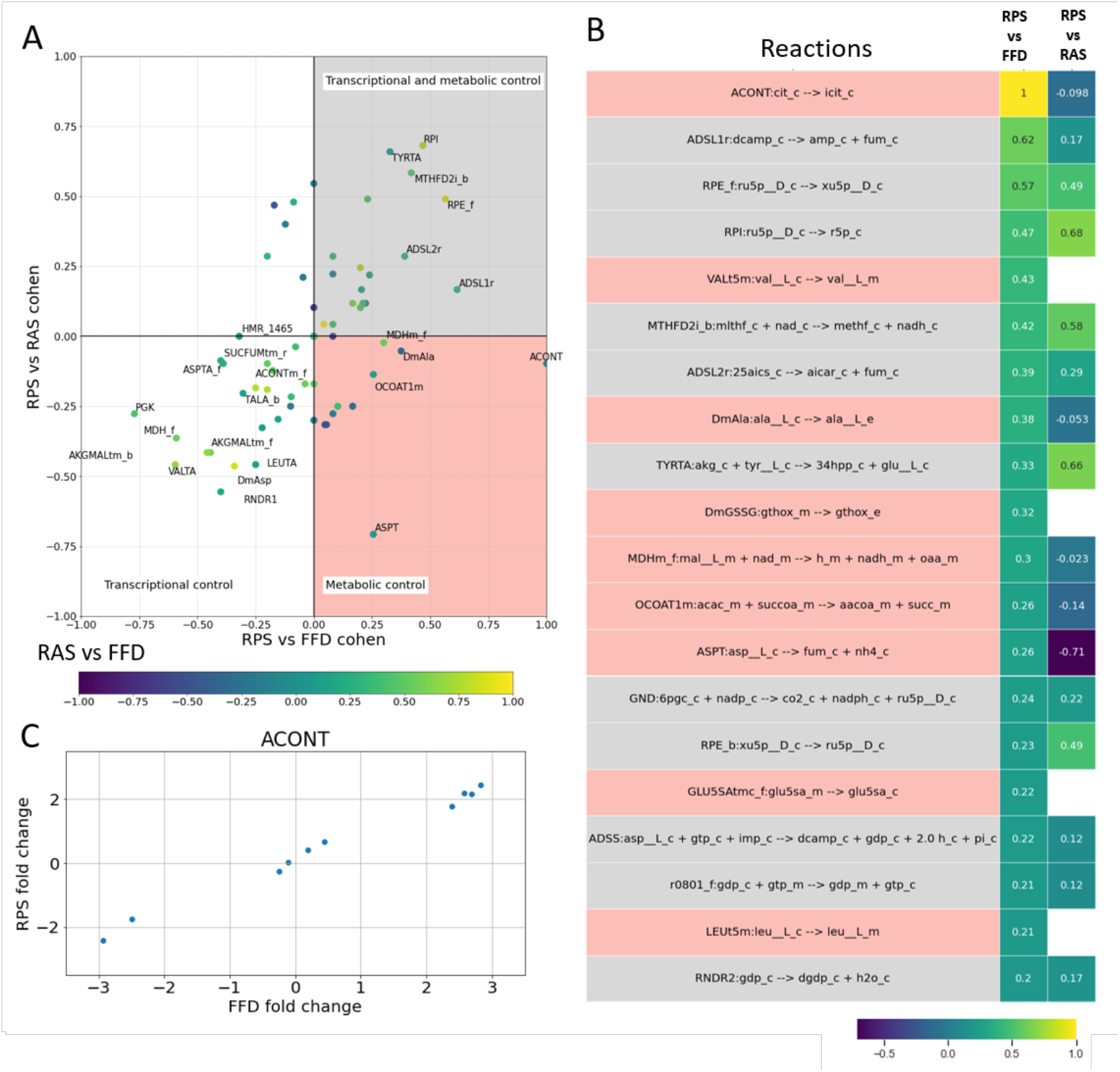
Variation concordance analysis. A) RPS vs FFD (x-axis) and the RPS vs RAS (y-axis) scores of the 81 metabolic reactions for which quantification of all substrate abundances was available. The points are coloured as a function the RAS vs FFD scores. We reported the names of the reactions having one of the scores greater than 0.2 at least (i.e. fair concordance). The points are coloured as a function the RAS vs FFD scores. B) Heatmap showing the RPS vs RAS and the RPS vs FFD concordance scores, for reactions having a level of concordance between RPS and FFD greater than 0.2. C) Scatterplot of average FFD log2-fold change vs average RPS log2-fold change for ACONT reaction.

The heatmap in Figure 4B reports the RPSvsRAS and the RPSvsFFD concordance scores (Cohen’s kappa) of ENGRO2 metabolic reactions, limited to the subset (of cardinality 81) of ENGRO2 reactions for which quantification of all substrate abundances was available. The values are ranked according to RPSvsFFD concordance scores. Only reactions with a RPSvsFFD concordance score higher than 0.2. Remaining reactions are reported in Supplementary Figure 3.

It can be observed that 11 reactions resulted from consistent transcriptional and metabolic regulation. On the contrary, 9 reactions resulted only metabolically regulated because of a RPSvsFFD score above 0.2 and a RPSvsRAS score below this threshold or even missing. Missing RPSvsRAS values occur when a reaction is not associated with a GPR. Among them, we obtained a perfect RPSvsFFD concordance of 1 for the ACONT reaction (Figure 4C), which catalyzes the production of cytosolic isocitrate from citrate.

### A few fluxes well discriminate the five cell lines

In Figure 5 the mean RPS (on the left) and the FFD (on the right) of the previously identified reactions resulting from a consistent transcriptional and metabolic regulation or only metabolically controlled are shown. For each reaction, the values are normalized by dividing them by the highest one.

**Fig 5.**
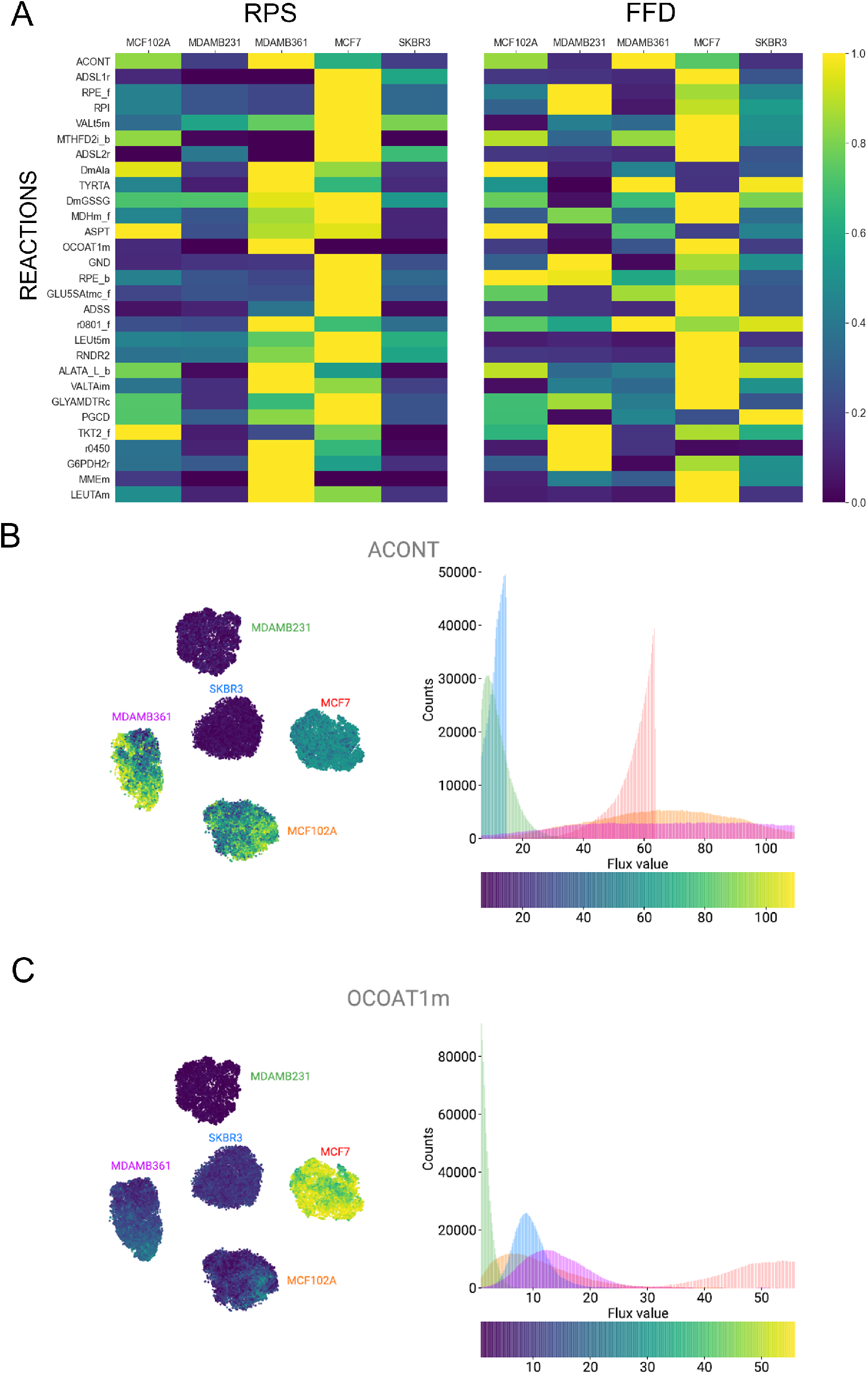
A) Normalized average RPS and median FFD for reactions in Figure 4B. B) Left: Figure 3A (constraints 1+2+3) with dots coloured according to the flux of cytosolic ACONT. Right: distribution of cytosolic ACONT flux values within the 5 cell lines. C) Same as B for mitochondrial OCOAT1m reaction.

We ranked FFDs according to their power in discriminating the five cell lines, as detailed in Material and Methods. The ranking is reported in Supplementary File 4.

The maps and histograms in Figure 5B and C report the FFD values distribution within the five cell lines of two metabolically regulated reactions that proved to discriminate the investigated cell lines: the ACONT reaction, which we previously discussed, and the mitochondrial OCOAT1m reaction catalysed by the 3-oxoacid CoA-transferase 1 (OXCT1), which transfers a CoA unit from succinyl-CoA to acetoacetate to form acetoacetyl-CoA and succinate. OXCT1 is a key enzyme involved in ketone bodies re-utilization by converting them into acetyl-CoA, which can enter the tricarboxylic acid cycle (TCA cycle), driving the production of ATP. Multiple studies elucidated the role of OXCT1 in breast cancer cells, proving to behave as a mitochondrial metabolic oncogene with a positive impact on the tumor growth and metastasis [46–48]. This evidence indicates ketone bodies inhibitors as a potential anti-tumor therapeutic strategy.

Among the metabolically regulated reactions identified in Figure 5, amino acid metabolism is heavily involved. Hence, we were interested in investigating whether amino acids are metabolized or synthesized by the network. To this aim, similarly to what was done in [32], we compared for each amino acid its uptake flux values against its contribution to biomass. The plots in Figure 6 report the results of this comparison for selected amino acids, namely isoleucine, valine, leucine and tyrosine are chosen. Sampled fluxes located along the bisector correspond to cases where all the internalized amino acid from the extracellular environment is wholly directed towards the production of biomass. This is the case of essential amino acid isoleucine (Figure 6A). Sampled fluxes located in the grey area above the bisector, as in the case of tyrosine (Figure 6D), indicate that the uptake rate of the amino acid is lower than its incorporation rate within biomass. This implies that it is synthesized by other metabolic processes within the network to fulfill the biomass requirement. On the contrary, when the sampled fluxes are below the bisector, as it is the case for valine and leucine (Figure 6B-C), the amino acid uptake rate is higher than its incorporation rate within biomass. This implies a dual role of the metabolite because in addition to contributing to biomass demand it is also metabolized within the network.

**Fig 6.**
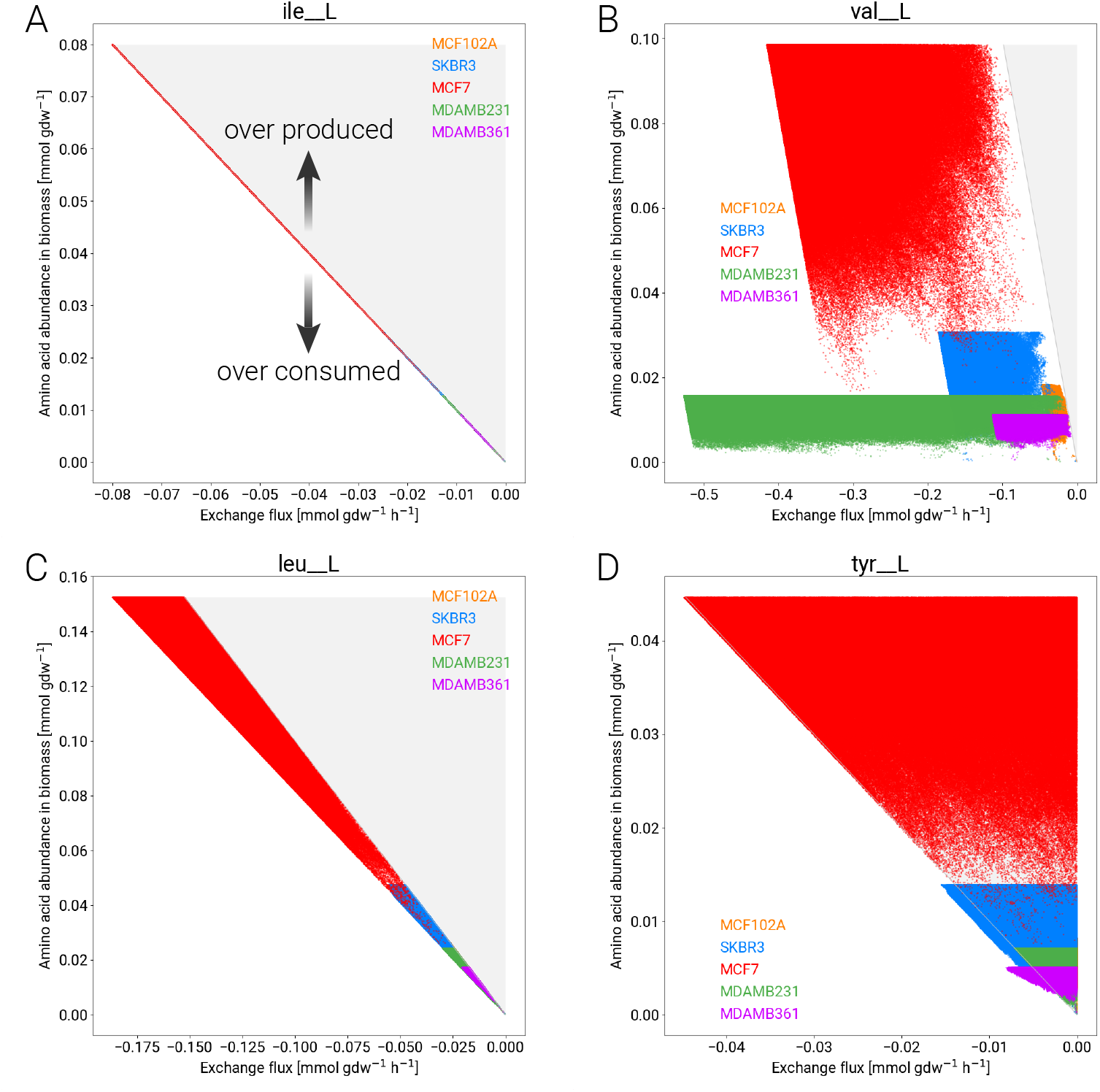
Amino acids requirement for growth when consumed from the extracellular environment and directed towards biomass synthesis. Each plot shows the sampled flux values of amino acid exchange reactions against their contribution to biomass production. Contribution to biomass is computed by multiplying the corresponding stoichiometric coefficient representing the amino acid abundance in biomass over the flux values of biomass reaction. Specifically, for each of the 1 milion sampled FFDs, the uptake flux of a given amino acid and the biomass flux were used for the calculation. Each panel refers to a different amino acids among those that were chosen as representative of alternative scenarios. A: isoleucine. B: valine. C: leucine. D: tyrosine.

Results in Figure 6 indicate that the five cell lines under study do not markedly differ in their preference to metabolize or synthesize a given amino-acid, but only in the extent at which it is metabolized or synthesized.

## Discussion

Metabolism is controlled by several interacting regulatory layers, including mechanisms that control the expression and activity of each metabolic enzyme and auto-regulatory features that depend on the interaction of metabolites with the enzymes [8, 49, 50]. Understanding whether a given reaction is controlled at the gene expression or at the metabolic level is required not only to understand regulatory features of metabolic pathways fully, but also to design appropriate actions to control metabolism in various fields, including health, wellness, and biotransformations [12], where metabolism plays a crucial regulatory role.

To enable this understanding we presented the INTEGRATE computational framework. INTEGRATE uses a metabolic stoichiometric model as a scaffold to predict metabolic flux distributions and their underlying regulation from transcriptomics and metabolomics data. In particular, we used ENGRO2, a manually curated, simulation-ready reconstruction of the human central carbon metabolism.

INTEGRATE first uses transcriptomic data to reconstruct metabolic fluxes, by constraining simulations of the ENGRO2 model. It then exploits intracellular metabolomic data to discriminate fluxes controlled at the substrate level from fluxes controlled by regulating enzyme activity or expression. To this aim, information on the stoichiometry of reactions included in the metabolic network model is used to compute Reaction Propensity Scores. INTEGRATE uses intracellular metabolomic datasets and the mass action law formulation to predict how differences in substrate availability translate into differences in metabolic fluxes (metabolic regulation only), neglecting enzymatic activity. Finally, INTEGRATE identifies fluxes that vary consistently at the metabolic and transcriptional regulation level and fluxes whose variation is concordant with metabolic regulation only.

INTEGRATE captures dynamic features of the metabolic state of different cells or tissues from the integration of high-throughput data that provide complementary views on their static profile. The model-based integration of transcriptomic and metabolomic data enriches their expressive power. Metabolic fluxes predicted by constraint-based modeling complement information on the differential activity of reactions derived from gene expression data, with the information on the direction of the observed variation.

Application of our pipeline to one non-tumorigenic cell line and four different breast cancer cell lines identified fluxes that markedly differ across the five cell lines, ascribing each differentially regulated flux to either transcriptional or metabolic control or both. Remarkably, we identified reactions for which there is a good agreement between flux variations and variations in RPSs. We remind that these two data sets are fully independent.

INTEGRATE also complements information on the differential propensity of reactions derived from metabolomics, with information on the compartment in which the reaction occurs. Metabolomic data alone would not have allowed us to differentiate between the contribution of the Aconitase reaction (ACONT) substrate when located in the cytosol compartment or when coming from its metabolic counterpart. On the contrary, INTEGRATE indicated that the metabolic regulation involves the cytosolic reaction. Notably, this information was complemented by transcriptomics data even though the aconitase flux itself is not regulated at the transcriptional level. This result indicates that indirect transcriptional regulation is likely to be responsible for observed differences in this flux, which would deserve further investigations.

The cytosolic ACONT reaction is strictly coupled to the cytosolic isocytrate dehydrogenase reaction, acting as an early participant in isocitrate dehydrogenase (IDH1)-dependent NADPH biosynthesis required for lipid biosynthesis [51]. Consistently, knockdown of the gene encoding cytosolic isociytrate dehydrogenase in preadipocytes results in decreased isocitrate dehydrogenase 1 (NADP+) mRNA levels and shifts the NADPH:NADP+ ratio towards the oxidized form, impairing adipogenesis [52]. These observation reinforce the involvement of cell redox balance in the metabolic rewiring of cancer metabolism, as already pointed out in [36, 53, 54].

Many analyses can be conceived and performed downstream of our pipeline. As an example, we analyzed fluxes that better discriminate between cell lines. We also investigated the metabolism of amino acids, to analyze whether they tend to be preferentially synthesized or metabolized by each cell line.

The direct exploitation of metabolomics represents the main novelty of our approach to determine whether a flux is regulated at the metabolic level. In a first instance, we intercept this information with information on transcriptional regulation. Nevertheless, the pipeline can be promptly extended to integrate proteomics and phosphoproteomics, allowing to characterize other hierarchical levels of regulation.

Knowing whether a flux is controlled at the metabolic or gene expression level is mandatory in designing therapeutic strategies. If a putative therapeutic flux is controlled metabolically, direct targeting the corresponding metabolic enzyme will not produce any effect. On the contrary, identifying the metabolic reaction(s) that indirectly affect the target reaction allows for designing an effective therapeutic intervention. Hence, by integrating high-throughput omics data through mathematical models, INTEGRATE makes it possible to dissect the complex and intertwined regulation of metabolic networks and inform targeted strategies to counteract metabolic rewiring and/or dysfunction underlying different pathological disorders.

The pipeline, however, can be applied to any case study. By way of example, metabolic engineering efforts are already inspired by constraint-based modeling [55]. Expanding the toolbox of computational tools will surely increase the success rate of efforts toward the application of engineering concepts to living organisms, contributing to the development of predictable, scalable, and efficient biological devices, whose performance is not hampered by inadequate knowledge of the underlying design principles [56].

## Material and Methods

### Model reconstruction

Throughout the process of model reconstruction, we evaluated the inclusion of a number of metabolic pathways according to their literature relevance in cancer metabolism.

In addition to the main changes introduced in ENGRO2 (as compared to ENGRO1) already enumerated in the Results Section, we included required cofactors within the stoichiometric equations of the model reactions. In fact, recent findings associated altered levels of some cofactors, including coenzyme A [57], water [58] and orthophosphate [59], with the emergence of various side effects and cancer promotion.

Moreover, to integrate gene expression data, we made all the reactions belonging to the oxidative phosphorylation (OXPHOS) pathway explicit. A lumped version of the entire route was instead considered in the ENGRO1 model in the form of two reactions explaining the NADH and FADH_2_ oxidation through the transfer of electrons from these two reducing agents to oxygen. The stoichiometry of the complex I-like reaction also included the generation of reactive oxygen species (ROS) that are known to be generated by a deficient Complex I activity following specific mutations affecting subunits of this enzymatic complex [60, 61]. In the ENGRO2 network, we included five separate reactions representing the reaction catalyzed by each OXPHOS complex. Because of the ROS production from the Complex I, it was necessary to add the ROS detoxification pathway by means of glutathione.

In view of the relevance of one-carbon metabolism in cancer cells [62], we enriched the ENGRO2 model by including both folate and the methionine cycles.

In ENGRO1, the oxidative branch of the pentose phosphate pathway was just partially included until the synthesis of the phosphoribosyl pyrophosphate (PRPP). PRPP is fundamental for cell biomass synthesis as the entry point of nucleotides biosynthesis. In view of the implications of the one-carbon metabolism in purines and pyrimidines production, we extended the pentose phosphate pathway by including in ENGRO2 the complete nucleotides biosynthesis route. Moreover, we also included all the non-oxidative branch of this pathway because of its relevance in reconverting the intermediates of this pathway into glycolytic metabolites.

Recent evidence highlighted a crucial role of beta oxidation in tumour cells for providing them fueling growth sources and benefiting tumour survival, especially under metabolic stress, such as following glucose or oxygen deprivation [63]. Moreover, fatty acid oxidation contributes to the total cell NADPH pool because of acetyl-CoA that, once produced, enters the TCA cycle and is converted with the oxaloacetate to citrate. The high relevance of NADPH is linked to its role in providing redox power for tumour cells to counteract the oxidative stress. For these reasons, we included the mitochondrial beta-oxidation pathway in ENGRO2 for degradating fatty acids that in the model are represented under the form of palmitate.

Another recent finding that we considered during the reconstruction of ENGRO2 regards the disregulation of the polyamines metabolism and its requirement under the neoplastic condition [64]. In view of these evidence, we added reactions belonging to the polyamines metabolism in the ENGRO2 model.

The biomass synthesis reaction was set according to biomass composition of Recon3D model in terms of metabolites and stoichiometric coefficient except for 1-Phosphatidyl-1D-Myo-Inositol, Phosphatidylcholine, Phosphatidylethanolamine, Phosphatidylglycerol, Cardiolipin, Phosphatidylserine, Sphingomyelin whose load is ascribed to palmitate.

### Flux Variabilty Analysis

Flux Variability Analysis (FVA) [65] is a constraint based modelling technique aimed at identifying the range of possible flux values for the reactions involved in the metabolic system under analysis. As other constraint based modelling methods, it relies on the fact that the steady states of the metabolic model corresponds to the kernel of the stoichiometric matrix *S* associated to the model itself. It is exactly this null space that defines the space of all the feasible fluxes. Moreover, to mimic as closely as possible the biological process in analysis, it is possible to bound the feasible fluxes space by means of convex half planes represented by the reaction’s lower and upper bounds.

Given a *M* × *N* stoichiometric matrix S, with M metabolites and N reactions, and the vectors ***ν_L_*** and ***ν_U_*** specifying, respectively, lower and upper bounds for the flux vector ***ν***, the mass balance constraint *S* · *ν* = 0 together with the flux boundaries specify the feasible region of the linear problem.

FVA solves the following two optimization problems (one for minimization and one for maximization) for each flux *ν_i_* of interest, with *i* = 1,…, *N*:

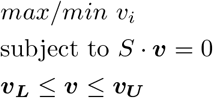

We remark that we do not ask the network to support a percentage of maximal possible biomass production rate as often done in other studies. We simply prevent biomass to be completely null, by setting the biomass synthesis lower bound to 10^-4^. This small level should be high enough to be distinguished form numeric instability.

### Reaction Activity Score computation

Reaction catalysis by a given set of enzymes is encoded within the model through the gene-protein-reaction (GPR) rule. GPR rules are logical expressions exploiting `AND` and `OR` logical operators to describe different types of relationship established among enzymes. In particular, `AND` operator is used when distinct genes encode multiple subunits of the same enzyme, implying that all the subunits are equally necessary for the reaction to take place. On the contrary `OR` operator is used when distinct genes encode multiple isoforms of the same enzyme, entailing that either isoform is enough for the reaction catalysis. These logical operators can be combined to describe more complex scenarios involving both isoforms and subunits.

According to [31], we combined the RNA-seq datasets with the defined GPR rules associated with each reaction *r* included in the model through the employment of the Reaction Activity Score (RAS). For each cell line *c* in the set *C* of the cell lines to analyze, for each sample *ξ* and for each reaction *r*, we computed the corresponding 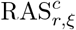, by resolving the corresponding logical expression taking the minimum transcript level value when multiple genes are joined by an `AND` operator, and taking the sum of their values when multiple genes joined by an `OR` operator encode distinct isoforms of the same enzyme.

We then computed the cell line reaction score by averaging over the samples as 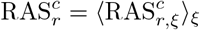. Once that 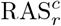 of each reaction is computed for each investigated cell line, these values are normalized on the maximum 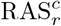 of all cell lines:

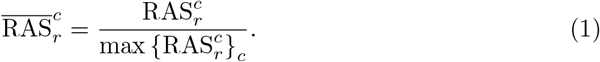

Finally, missing information about transcript level implies that RAS of the corresponding reactions are set to 1. When RAS is equal to 0 in all the cell line, the corresponding normalized 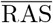 is kept equal to 0 to consider that the corresponding reaction is off in all the cell lines.

### Cell-relative model construction

#### Constraints on nutrient availability

To set constraints on nutrient availability, we defined the set of nutrients that can be internalized by each of the five investigated cell lines according to the two exploited experimental medium composition. For every uptaken metabolite, an exchange reaction is included within the network by setting its upper bound to 0 and tuning its lower bound proportionally to the corresponding concentration contained in the growth medium.

#### Constraints on extracellular fluxes

To set constraints on the extracellular fluxes according to the experimentally determined flux ratio of lactate to glucose, lactate to glutamine and glutamate to glutamine, we proceeded as follows:

- Firstly, we took the concentration values of the two nutrients glucose, glutamine and of the two byproducts glutamate and lactate previously collected from the YSI analyzer of spent medium.
- Treating the two biological replicas separately, we computed for each of the three technical replicas the mean concentration difference of the considered metabolites between initial time and after 48 hours of growth to get their utilization or formation rate. Focusing on the ratio between these metabolites rather than on their absolute uptake or secretion rates, we limited to consider the mean concentration difference values between time 0 h and 48 h without dividing them by the integral of cells number.
- We computed the lactate produced over glucose consumed, lactate produced over glutamine consumed and glutamate produced over glutamine consumed ratios for the two biological replicas.
- We then added these experimentally determined flux ratios as further constraints to the model to ensure that exchange reactions involved in the considered ratios follow the experimental values. To give an example, the lactate to glucose ratio is included within the model as constraint by considering the following expression:

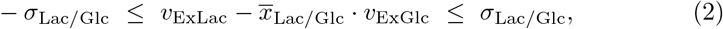

where the ratio of the flux value *ν*_ExLac_ of the lactate secretion reaction over the flux value *ν*_ExGlc_ of the glucose consumption reaction ranges between minus one and plus one standard deviation *σ*_*Lac/Glc*_ of the mean lactate to glucose ratio 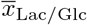 of the two biological replicas.

#### Trascriptomics-derived constraints

Assuming that it is possible to modulate the reactions activity by transcription mechanisms, it is possible to exploit the 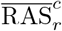 score to further constraint the extremal fluxes, ***ν_L_***, ***ν_U_*** obtained through FVA, to represent this process and better mimic the cell lines behaviour:

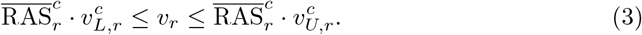

In this case, for the internal reactions when a GPR rule exists for a given reaction, its lower and upper bounds are scaled according to its computed 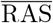.

The only exceptions to application of this constraints are the boundaries for reactions CARPEPT1tc and HIStiDF for which we ignored the 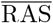 contribution because the constraints were limiting too severely the growth of some of the cellular lines.

### Generation of FFD dataset, via random sampling

Once the cell-relative models were created, for following analyses, we converted them into irreversible models, in which reversible reactions are represented with two distinct and complementary forward reactions.

The ability to properly sample the constrained null space of *S* is of paramount importance to obtain correct fluxes distributions.

In this work, we exploited the implementation of optGpSampler algorithm [66] available in COBRApy [67], and we sampled a million of steady-state solutions of the ENGRO2 model in all the tested conditions.

### Generation of RPS dataset

Let *x^i^* be the vector of abundances of the chemical species in a given steady state *i* of the metabolic network. Let *ν_i_* be the vector of reaction flux rates in the same steady state *i*. The flux of chemical species through a reaction is the rate of the forward reaction, minus that of the reverse reaction (in molecules per unit of time). When dealing with irreversible models, the reaction flux and the rate coincide.

The following assumptions allow one to analytically estimate relative fluxes from relative abundances:

- for each reaction *r* in the system, the mass action law is assumed: 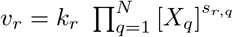, where *k_r_* is the kinetic constant of reaction *r* and *X_q_* is the *q^th^* substrate of the *N* total substrates of reaction *r*, and *s*_*r*,*q*_ is the stoichiometric coefficient of substrate *X_q_* in reaction *r* i.e., how many molecules of the substrate partake to the reaction;
- the kinetic constant *k_r_* of a given reaction *r* is assumed to not vary between two steady states *i* and *j*.

Given such assumptions, the variation between the flux of an irreversible reaction *r* in two steady states *i* and *j* can thus be computed as the ratio 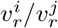:

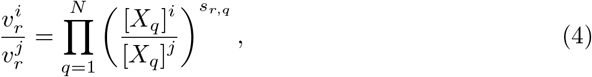

which does not depend on *k_r_*.

It goes without saying that if the numerator is higher than the denominator, than flux *ν_r_* in steady state *i* is higher than flux *ν_r_* in steady state *j*. Therefore, in order to compare the susbtrate contribution to the reaction rate in different cell lines, we computed for each reaction *r* and for each cell line *c* (assumed at steady state) a Reaction Propensity Score (RPS), defined as as follows:

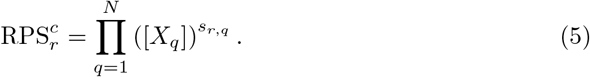

Of course, the above assumptions not often hold. However, RPS_*r*_ allow us to predict whether the flux of a reaction in a cell line is expected to be different as compared to another cell line, just considering substrate availability. Nothing prevents the observed flux variation to be consistent with the RPS variation, even if the enzyme activity actually differs between the two cell lines under study.

### Concordance analysis

We wanted to test whether the variation in the value of a reaction is consistent between different datasets. Statistical evaluation of the concordance between two techniques used to measure a particular variable, under identical circumstances, has been largely addressed in literature [68]. We decided to treat our measurements, i.e. the log ratio of the flux, as up, down or no-change.

Concerning the FFD values, we performed the Mann-Whitney U test [69] (p-value < 0.05) between the distributions of each pair of the five investigated cell lines to determine if the FFD of every reaction differed significantly between the two cell lines. In parallel, the log_2_ of the absolute ratio of the median flux of each reaction in the two cell lines is also computed.

Concerning the RAS and the RPS values, we performed a t-test (p-value < 0.05) for the means of each pair of RAS and RPS samples taken for all the pairwise combinations of the five investigated cell lines to determine if the values differed significantly between the two cell lines. We also computed the log_2_ of the absolute ratio of the mean RAS and RPS values of each reaction in the two cell lines.

Hence, we registered the sign of the variation for each pair of the five cell lines under study (for a total of 10 pairs) according to each of the three datasets. At first instance, a positive sign is registered if the distribution of samples values of first member of the comparison is statistically higher and if the average or median value is at least 20% higher. A negative sign is registered if the distribution of samples values of the first member of the comparison is statistically lower and if the average or median value is at least 20% lower. A 0 is registered otherwise. We used a relaxed threshold for the fold-change as in [31] because even a difference of 20% in genes encoding members of a metabolic pathway may dramatically alter the flux through the pathway. Yet, this parameter can be modified arbitrarily.

We quantified the level of concordance of the 10 variation signs (1 for each pair of cell lines) for a given pair of datasets by means of the Cohen’s kappa metric, which has been commonly used to measure inter-rater reliability for qualitative (categorical) items.

### Ranking of discriminative fluxes

For each reaction *r* in our model, we tried to address the problem of quantifying its capability to fingerprint the investigated cell lines. To this aim, we derived the Reaction Fingerprinting Score (RFS), which exploits the frequency distribution of flux values derived from the optGpSampler algorithm and applies a homogeneous binning among the cell lines (i.e. Rice’s rules). According to this, the frequency distribution 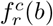 retains the flux values count for cell line *c* of reaction *r* in the binning *b*:

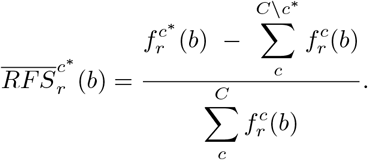

To highlight the fingerprinting role of reaction *r*, we chose to keep into account only the contribution to counts for which cell line *c*\* is greater than the contribution of other cell lines *C*\*c*\*:

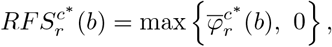

whereas with 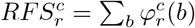 it is possible to identify cell lines with a more set apart fluxes distribution with respect to the others cell lines. The same 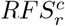 allows to state which reactions are more discriminating to fingerprint cell line *c*. With the aim to identify those reactions that more than others allow to globally discriminate the set of cell lines in analysis, it is necessary to combine this information defining the RFS as

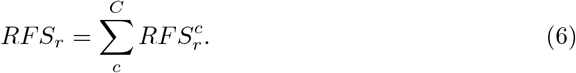

### Cell culture

MDA-MB231 and MCF7 cell lines were grown in Dulbecco’s modified Eagle’s medium (DMEM) containing 4 mM L-glutamine, supplemented with 10% fetal bovine serum (FBS). MDA-MB361 cell line was maintained in DMEM/F12 containing 20% FBS and 4 mM L-glutamine. SKBR3 cell line was grown in DMEM containing 2 mM L-glutamine, supplemented with 10% FBS. MCF102A cell line was maintained in DMEM/F-12 containing 5% horse serum, 2.5 mM L-glutamine, 20 ng/ml EGF, 100 ng/ml cholera toxin, 0.01 mg/ml insulin, and 500 ng/ml hydrocortisone. All media were supplemented with 100 U/ml penicillin and 100 μg/ml streptomycin, and cells were incubated at 37°C in a 5% CO2 incubator. All reagents for media were purchased from Life Technologies (Carlsbad, CA, USA).

### Cell proliferation and protein content analysis

Cells were plated in 6-well plates in normal growth medium. Culture medium was replaced after 18 h and cells were collected and counted after 24, 48, and 72 hours. Protein extraction was performed at each indicated time in RIPA buffer supplemented with protease inhibitor cocktail. Protein content was evaluated through Bradford assay, using Pierce™ Coomassie Plus (Thermo Scientific) and bovine serum albumin as a standard protein.

### Metabolite extraction from cell culture

Cells were plated in 6-well plates with normal growth medium. The culture medium was replaced after 18 h and then cells were incubated for 48 h. For metabolites extraction, cells were quickly rinsed with NaCl 0.9% and quenched with 500 *μ*l ice-cold 70:30 acetonitrile-water. Plates were placed at −80°C for 10 minutes, then the cells were collected by scraping, sonicated 5 seconds for 5 pulses at 70% power twice and then centrifuged at 12000 g for 10 min at 4°C. The supernatant was collected in a glass insert and evaporated in a centrifugal vacuum concentrator (Concentrator plus/ Vacufuge^®^ plus, Eppendorf) at 30 °C for about 2.5 h. Samples were then resuspended with 150 ^μ^l of ultrapure water prior to analyses.

### LC-MS metabolic profiling

LC separation was performed using an Agilent 1290 Infinity UHPLC system and an InfintyLab Poroshell 120 PFP column (2.1 x 100 mm, 2.7 μm; Agilent Technologies). The injection volume was 15 μL, the flow rate was 0.2 mL/min with column temperature set at 35°C. Both mobile phase A (100% water) and B (100% acetonitrile) contained 0.1% formic acid. LC gradient conditions were: 0 min: 100% A; 2 min: 100% A; 4 min: 99% A; 10 min: 98% A; 11 min: 70% A; 15 min: 70% A; 16 min: 100% A with 5 min of post-run. MS detection was performed using an Agilent 6550 iFunnel Q-TOF mass spectrometer with Dual JetStream source operating in negative ionization mode. MS parameters were: gas temp: 285°C; gas flow: 14 L/min; nebulizer pressure: 45 psig; sheath gas temp: 330°C; sheath gas flow: 12 L/min; VCap: 3700 V; Fragmentor: 175 V; Skimmer: 65 V; Octopole RF: 750 V. Active reference mass correction was done through a second nebulizer using the reference solution (m/z 112.9855 and 1033.9881) dissolved in the mobile phase 2-propanol-acetonitrile-water (70:20:10 v/v). Data were acquired from m/z 60–1050. Data analysis and isotopic natural abundance correction were performed with MassHunter ProFinder (Agilent).

### Metabolites quantification in the media samples

Absolute quantification of glucose, lactate, glutamine, and glutamate in spent media after 48 hours of growth was determined enzymatically using YSI2950 bioanalyzer (YSI Incorporated, Yellow Springs, OH, USA).

### RNA Extraction

Total RNA was extracted from at least 8 x 106 cells by using RNeasy Mini Kit (Qiagen). Each pellet was resuspended by adding 30 *μ*L of RNAse-free water. Following RNA isolation, DNAse treatment was performed using DNAse I, RNase-free (ThermoFisher Scientific). After purification, to assess the final RNA yield and purity, the spectrophotometer NanoDrop ND-1000 (NanoDrop Technologies) was employed by the means of 260/280 and 260/230 ratios. The 2200 TapeStation instrument (Agilent Technologies) also evaluated the RNA quality, in order to assess the RNA Integrity Number Equivalent (RIN) for each processed sample.

### RNA-Seq library preparation

The RNA-Seq libraries were prepared using the Illumina TruSeq Stranded mRNA Library Prep Kit, according to the manufacturer’s instructions. Starting from each RNA isolated, three replicates were prepared. All the 15 obtained libraries were, as first, analyzed by the 2200 TapeStation instrument to check their length and quality and, then, quantified by the fluorescent dye PicoGreen^®^ (ThermoFisher Scientific) on NanoDrop ND-1000 to calculate the concentration. RNA-Seq libraries were then diluted at 2 nM concentration and normalized using standard library quantification and quality control procedures as recommended by the Illumina protocol. RNA-Seq libraries were sequenced using Illumina^®^ HiSeq2500 to obtain 150-bp paired-end reads. After fastq quality control by using FastQC tool, raw reads were mapped with STAR aligner (v.2.6.1d) to human reference genome (hg38) and gene counts were calculated by HTSeq (v.0.6.1), using the hg38 Encode-Gencode GTF file (v28) as gene annotation file. Gene abundance was measured in fragments per kb of exon per million fragments mapped (FPKM).

## Acknowledgments

The institutional financial support to SYSBIO.ISBE.IT within the Italian Roadmap for ESFRI Research Infrastructures and the FLAG-ERA grant ITFoC are gratefully acknowledged. Financial support from the Italian Ministry of University and Research (MIUR) through grant ‘Dipartimenti di Eccellenza 2017’ to University of Milano Bicocca is also greatly acknowledged. We warmly thank Alex Graudenzi for useful suggestions.

## Author contributions

Conceptualization: CD, LA, MV Data Curation: BGG, MDF Formal Analysis: DP, MDF Funding Acquisition: MV, LA Investigation: CD, DP, MDF, DG, MB, EM, CC Methodology: CD, DP, MDP Software: MDP, CD, BGG Supervision: CD Visualization: CD, MDF, DP, BGG Writing – Original Draft Preparation: CD, MDP Writing – Review and Editing: MV, LA, DP

## Supporting information

**S1 Fig. ENGRO2 map - part I.** Graphical representations of central carbon metabolism reactions included in ENGRO2 model.

**S2 Fig. ENGRO2 map - part II.** Graphical representations of essential amino acid metabolism reactions included in ENGRO2 model.

**S3 Fig. Concordance scores.** Heatmap of RPSvsRAS and RPSvsFFD Cohen’s kappa coefficients for reactions not reported in Figure 4.

**S1 File. Input experimental datasets.** Read counts (FPKM), LC-MS metabolic profiling and extracellular fluxes.

**S2 File. ENGRO2 model.** SBML of the unconstrained model.

**S3 File. Cell relative metabolic models.** Compressed file archive including the five cell-relative models (SBML).

**S4 File. Flux ranking.** Engro2 model reactions scored according to the distance of the FFDs distributions of the five cell lines.

## References

1. Nielsen J. Systems biology of metabolism: a driver for developing personalized and precision medicine. Cell metabolism. 2017;25(3):572–579.

2. Procaccini C, Santopaolo M, Faicchia D, Colamatteo A, Formisano L, De Candia P, et al. Role of metabolism in neurodegenerative disorders. Metabolism. 2016;65(9):1376–1390.

3. Noda-Garcia L, Liebermeister W, Tawfik DS. Metabolite–enzyme coevolution: from single enzymes to metabolic pathways and networks. Annual Review of Biochemistry. 2018;87:187–216.

4. Noor E, Flamholz A, Bar-Even A, Davidi D, Milo R, Liebermeister W. The protein cost of metabolic fluxes: prediction from enzymatic rate laws and cost minimization. PLoS computational biology. 2016;12(11):e1005167.

5. He F, Stumpf MP. Quantifying dynamic regulation in metabolic pathways with nonparametric flux inference. Biophysical journal. 2019;116(10):2035–2046.

6. Rossell S, van der Weijden CC, Lindenbergh A, van Tuijl A, Francke C, Bakker BM, et al. Unraveling the complexity of flux regulation: a new method demonstrated for nutrient starvation in Saccharomyces cerevisiae. Proceedings of the National Academy of Sciences. 2006;103(7):2166–2171.

7. Millard P, Smallbone K, Mendes P. Metabolic regulation is sufficient for global and robust coordination of glucose uptake, catabolism, energy production and growth in Escherichia coli. PLoS computational biology. 2017;13(2):e1005396.

8. Millard P, Smallbone K, Mendes P. Metabolic regulation is sufficient for global and robust coordination of glucose uptake, catabolism, energy production and growth in Escherichia coli. PLoS computational biology. 2017;13(2):e1005396.

9. Reid MA, Dai Z, Locasale JW. The impact of cellular metabolism on chromatin dynamics and epigenetics. Nature cell biology. 2017;19(11):1298–1306.

10. Wang YP, Lei QY. Metabolite sensing and signaling in cell metabolism. Signal transduction and targeted therapy. 2018;3(1):1–9.

11. Lempp M, Farke N, Kuntz M, Freibert SA, Lill R, Link H. Systematic identification of metabolites controlling gene expression in E. coli. Nature communications. 2019;10(1):1–9.

12. Damiani C, Gaglio D, Sacco E, Alberghina L, Vanoni M. Systems metabolomics: From metabolomic snapshots to design principles. Current opinion in biotechnology. 2020;63:190–199.

13. Cascante M, Marin S. Metabolomics and fluxomics approaches. Essays in biochemistry. 2008;45:67.

14. Allen DK, Young JD. Tracing metabolic flux through time and space with isotope labeling experiments. Current opinion in biotechnology. 2020;64:92–100.

15. Hirai MY, Yano M, Goodenowe DB, Kanaya S, Kimura T, Awazuhara M, et al. Integration of transcriptomics and metabolomics for understanding of global responses to nutritional stresses in Arabidopsis thaliana. Proceedings of the National Academy of Sciences. 2004;101(27):10205–10210.

16. Liu P, Luo J, Zheng Q, Chen Q, Zhai N, Xu S, et al. Integrating transcriptome and metabolome reveals molecular networks involved in genetic and environmental variation in tobacco. DNA Research. 2020;27(2):dsaa006.

17. Hassan MA, Al-Sakkaf K, Shait Mohammed MR, Dallol A, Al-Maghrabi J, Aldahlawi A, et al. Integration of transcriptome and metabolome provides unique insights to pathways associated with obese breast cancer patients. Frontiers in oncology. 2020;10:804.

18. Ren S, Shao Y, Zhao X, Hong CS, Wang F, Lu X, et al. Integration of metabolomics and transcriptomics reveals major metabolic pathways and potential biomarker involved in prostate cancer. Molecular & Cellular Proteomics. 2016;15(1):154–163.

19. Zimmermann M, Kogadeeva M, Gengenbacher M, McEwen G, Mollenkopf HJ, Zamboni N, et al. Integration of metabolomics and transcriptomics reveals a complex diet of Mycobacterium tuberculosis during early macrophage infection. MSystems. 2017;2(4).

20. Zhang L, Ma C, Chao H, Long Y, Wu J, Li Z, et al. Integration of metabolome and transcriptome reveals flavonoid accumulation in the intergeneric hybrid between Brassica rapa and Raphanus sativus. Scientific reports. 2019;9(1):1–8.

21. Cavill R, Jennen D, Kleinjans J, Briedé JJ. Transcriptomic and metabolomic data integration. Briefings in bioinformatics. 2016;17(5):891–901.

22. Siddiqui JK, Baskin E, Liu M, Cantemir-Stone CZ, Zhang B, Bonneville R, et al. IntLIM: integration using linear models of metabolomics and gene expression data. BMC bioinformatics. 2018;19(1):1–12.

23. Opdam S, Richelle A, Kellman B, Li S, Zielinski DC, Lewis NE. A systematic evaluation of methods for tailoring genome-scale metabolic models. Cell systems. 2017;4(3):318–329.

24. Machado D, Herrgård M. Systematic evaluation of methods for integration of transcriptomic data into constraint-based models of metabolism. PLoS Comput Biol. 2014;10(4):e1003580.

25. Jamialahmadi O, Hashemi-Najafabadi S, Motamedian E, Romeo S, Bagheri F. A benchmark-driven approach to reconstruct metabolic networks for studying cancer metabolism. PLoS computational biology. 2019;15(4):e1006936.

26. Sajitz-Hermstein M, Töpfer N, Kleessen S, Fernie AR, Nikoloski Z. iReMet-flux: constraint-based approach for integrating relative metabolite levels into a stoichiometric metabolic models. Bioinformatics. 2016;32(17):i755–i762.

27. Pandey V, Hadadi N, Hatzimanikatis V. Enhanced flux prediction by integrating relative expression and relative metabolite abundance into thermodynamically consistent metabolic models. PLoS computational biology. 2019;15(5):e1007036.

28. Yizhak K, Benyamini T, Liebermeister W, Ruppin E, Shlomi T. Integrating quantitative proteomics and metabolomics with a genome-scale metabolic network model. Bioinformatics. 2010;26(12):i255–i260.

29. Cakir T, Patil KR, Önsan ZI, Ülgen KÖ, Kirdar B, Nielsen J. Integration of metabolome data with metabolic networks reveals reporter reactions. Molecular systems biology. 2006;2(1):50.

30. Katzir R, Polat IH, Harel M, Katz S, Foguet C, Selivanov VA, et al. The landscape of tiered regulation of breast cancer cell metabolism. Scientific reports. 2019;9(1):1–12.

31. Graudenzi A, Maspero D, Di Filippo M, Gnugnoli M, Isella C, Mauri G, et al. Integration of transcriptomic data and metabolic networks in cancer samples reveals highly significant prognostic power. Journal of Biomedical Informatics. 2018;87:37–149.

32. Nilsson A, Haanstra JR, Engqvist M, Gerding A, Bakker BM, Klingmüller U, et al. Quantitative analysis of amino acid metabolism in liver cancer links glutamate excretion to nucleotide synthesis. Proceedings of the National Academy of Sciences. 2020;117(19):10294–10304.

33. Sánchez BJ, Zhang C, Nilsson A, Lahtvee PJ, Kerkhoven EJ, Nielsen J. Improving the phenotype predictions of a yeast genome-scale metabolic model by incorporating enzymatic constraints. Molecular systems biology. 2017;13(8):935.

34. Brunk E, Sahoo S, Zielinski DC, Altunkaya A, Dräger A, Mih N, et al. Recon3D enables a three-dimensional view of gene variation in human metabolism. Nature biotechnology. 2018;36(3):272.

35. Nobile MS, Coelho V, Pescini D, Damiani C. Accelerated global sensitivity analysis of genome-wide constraint-based metabolic models. BMC bioinformatics. 2021;22(2):1–17.

36. Damiani C, Colombo R, Gaglio D, Mastroianni F, Pescini D, Westerhoff HV, et al. A metabolic core model elucidates how enhanced utilization of glucose and glutamine, with enhanced glutamine-dependent lactate production, promotes cancer cell growth: The WarburQ effect. PLoS computational biology. 2017;13(9):e1005758.

37. Di Filippo M, Colombo R, Damiani C, Pescini D, Gaglio D, Vanoni M, et al. Zooming-in on cancer metabolic rewiring with tissue specific constraint-based models. Computational biology and chemistry. 2016;62:60–69.

38. Lytovchenko O, Kunji ER. Expression and putative role of mitochondrial transport proteins in cancer. Biochimica et Biophysica Acta (BBA)-Bioenergetics. 2017;1858(8):641–654.

39. Ward PS, Thompson CB. Metabolic reprogramming: a cancer hallmark even warburg did not anticipate. Cancer cell. 2012;21(3):297–308.

40. Flaveny CA, Griffett K, El-Gendy BEDM, Kazantzis M, Sengupta M, Amelio AL, et al. Broad anti-tumor activity of a small molecule that selectively targets the Warburg effect and lipogenesis. Cancer cell. 2015;28(1):42–56.

41. Patra S, Ghosh A, Roy SS, Bera S, Das M, Talukdar D, et al. A short review on creatine–creatine kinase system in relation to cancer and some experimental results on creatine as adjuvant in cancer therapy. Amino Acids. 2012;42(6):2319–2330.

42. Rabinovich S, Adler L, Yizhak K, Sarver A, Silberman A, Agron S, et al. Diversion of aspartate in ASS1-deficient tumours fosters de novo pyrimidine synthesis. Nature. 2015;527(7578):379.

43. Weglarz-Tomczak E, Mondeel TD, Piebes DG, Westerhoff HV. Simultaneous integration of gene expression and nutrient availability for studying the metabolism of hepatocellular carcinoma cell lines. Biomolecules. 2021;11(4):490.

44. Van der Maaten L, Hinton G. Visualizing data using t-SNE. Journal of machine learning research. 2008;9(11).

45. Ashby D. Practical statistics for medical research. Douglas G. Altman, Chapman and Hall, London, 1991. No. of pages: 611. Price:£3.00; 1991.

46. Ozsvari B, Sotgia F, Simmons K, Trowbridge R, Foster R, Lisanti MP. Mitoketoscins: novel mitochondrial inhibitors for targeting ketone metabolism in cancer stem cells (CSCs). Oncotarget. 2017;8(45):78340.

47. Zhang S, Xie C. The role of OXCT1 in the pathogenesis of cancer as a rate-limiting enzyme of ketone body metabolism. Life sciences. 2017;183:110–115.

48. Martinez-Outschoorn UE, Lin Z, Whitaker-Menezes D, Howell A, Sotgia F, Lisanti MP. Ketone body utilization drives tumor growth and metastasis. Cell cycle. 2012;11(21):3964–3971.

49. Gonçalves E, Raguz Nakic Z, Zampieri M, Wagih O, Ochoa D, Sauer U, et al. Systematic analysis of transcriptional and post-transcriptional regulation of metabolism in yeast. PLoS computational biology. 2017;13(1):e1005297.

50. Ortmayr K, Dubuis S, Zampieri M. Metabolic profiling of cancer cells reveals genome-wide crosstalk between transcriptional regulators and metabolism. Nature communications. 2019;10(1):1–13.

51. Koh HJ, Lee SM, Son BG, Lee SH, Ryoo ZY, Chang KT, et al. Cytosolic NADP+-dependent isocitrate dehydrogenase plays a key role in lipid metabolism. Journal of Biological Chemistry. 2004;279(38):39968–39974.

52. Moreno M, Ortega F, Xifra G, Ricart W, Fernandez-Real JM, Moreno-Navarrete JM. Cytosolic aconitase activity sustains adipogenic capacity of adipose tissue connecting iron metabolism and adipogenesis. The FASEB Journal. 2015;29(4):1529–1539.

53. Alberghina L, Gaglio D. Redox control of glutamine utilization in cancer. Cell death & disease. 2014;5(12):e1561–e1561.

54. Lewis JE, Forshaw TE, Boothman DA, Furdui CM, Kemp ML. Personalized genome-scale metabolic models identify targets of redox metabolism in radiation-resistant tumors. Cell Systems. 2021;12(1):68–81.

55. Nielsen J, Keasling JD. Engineering cellular metabolism. Cell. 2016;164(6):1185–1197.

56. Kwok R. Five hard truths for synthetic biology. Nature News. 2010;463(7279):288–290.

57. Srinivasan B, Sibon OC. Coenzyme A, more than ‘just’a metabolic cofactor; 2014.

58. Papadopoulos MC, Saadoun S. Key roles of aquaporins in tumor biology. Biochimica et Biophysica Acta (BBA)-Biomembranes. 2015;1848(10):2576–2583.

59. Sapio L, Naviglio S. Inorganic phosphate in the development and treatment of cancer: A Janus Bifrons? World journal of clinical oncology. 2015;6(6):198.

60. Urra FA, Muñoz F, Lovy A, Cárdenas C. The mitochondrial complex (I) ty of cancer. Frontiers in oncology. 2017;7:118.

61. Liou GY, Storz P. Reactive oxygen species in cancer. Free radical research. 2010;44(5):479–496.

62. Newman AC, Maddocks OD. One-carbon metabolism in cancer. British journal of cancer. 2017;116(12):1499.

63. Qu Q, Zeng F, Liu X, Wang Q, Deng F. Fatty acid oxidation and carnitine palmitoyltransferase I: emerging therapeutic targets in cancer. Cell death & disease. 2017;7(5):e2226.

64. Murray-Stewart TR, Woster PM, Casero RA. Targeting polyamine metabolism for cancer therapy and prevention. Biochemical Journal. 2016;473(19):2937–2953.

65. Gudmundsson S, Thiele I. Computationally efficient flux variability analysis. BMC bioinformatics. 2010;11(1):1–3.

66. Megchelenbrink W, Huynen M, Marchiori E. optGpSampler: an improved tool for uniformly sampling the solution-space of genome-scale metabolic networks. PloS one. 2014;9(2):e86587.

67. Ebrahim A, Lerman JA, Palsson BO, Hyduke DR. COBRApy: constraints-based reconstruction and analysis for python. BMC systems biology. 2013;7(1):1–6.

68. Kwiecien R, Kopp-Schneider A, Blettner M. Concordance analysis: part 16 of a series on evaluation of scientific publications. Deutsches Arzteblatt International. 2011;108(30):515.

69. Mann HB, Whitney DR. On a test of whether one of two random variables is stochastically larger than the other. The annals of mathematical statistics. 1947; p. 50–60.

